# Tumor MHCII immunity requires in situ antigen presentation by cancer-associated fibroblasts

**DOI:** 10.1101/2020.03.24.005355

**Authors:** Dimitra Kerdidani, Emmanouil Aerakis, Kleio-Maria Verrou, Petros Stamoulis, Katerina Goudevenou, Alejandro Prados, Christos Tzaferis, Ioannis Vamvakaris, Evangelos Kaniaris, Konstantinos Vachlas, Evangelos Sepsas, Konstantinos Potaris, Anastasios Koutsopoulos, Maria Tsoumakidou

## Abstract

A key unknown of the functional space in tumor immunity is whether physiologically relevant cancer antigen presentation occurs solely in draining lymph nodes versus tumors. Professional antigen presenting cells, i.e. the dendritic cells, are scarce and immature within tumors, greatly outnumbered by MHCII expressing non-hematopoietic cells, such as antigen-presenting cancer-associated fibroblasts (apCAFs). We hypothesized that after their exit from tumor-draining lymph nodes T cells depend on a second wave of antigen presentation provided in situ by structural cells. We show that dense apCAF regions in human lung tumors define hot immunological spots with increased numbers of CD4 T cells. The transcriptomic profile of human lung apCAFs aligned to that of pancreatic apCAFs across mice and humans and were both enriched for alveolar type II genes, suggesting an epithelial origin. Mechanistically, human apCAFs directly activated the TCRs of adjacent effector CD4 T cells and at the same time produced high levels of c1q, which acted on surface c1qbp on T cells to rescue them from apoptosis. Fibroblast-specific deletion of MHCII in mice impaired local MHCII immunity and accelerated tumor growth, while inducing c1qbp overexpression in adoptively transferred T cells expanded their numbers within tumors and reduced tumour burden. Collectively, our work shows that tumor T cell immunity post lymph node exit requires peripheral antigen presentation by a subset of CAFs and proposes a new conceptual framework upon which effective cancer immunotherapies can be built.

## INTRODUCTION

The series of immunological events that takes place between tumors and tumor draining lymph nodes forms a cyclic trajectory that is being referred to as the cancer-immunity cycle (1). In the first step of these events tumor antigens are carried to the tumor draining lymph nodes (LNs) and partly transferred to resident dendritic cells (DCs) (2). In LNs, migratory and resident DCs present their antigenic cargo to antigen-inexperienced (naïve) T cells, which become differentiated effector cells that egress from LNs and enter tumors. In tumors, CD8 cells exhibit direct killing activity against cancer cells, but they are seriously dependent on CD4 T cells for function and transition to memory cells (3–6). Although our current understanding of the functional space in the cancer-immunity cycle is that cancer antigen presentation primarily occurs in lymph nodes, the contribution of in situ cancer antigen presentation has not been be ruled out (7, 8).

Very few studies have directly addressed the role of peripheral antigen presentation in T cell responses (7, 9–12). In cancer three lines of evidence support that the TCRs are stimulated in situ within solid tumors. First, the CD4 TCR repertoire is regionally shaped by the local neoantigen load (13). Second, stem-like CD8 T cells reside in dense MHCII expressing cell niches within tumors (14). Third, right flank tumours that differ only in one MHCII neoantigen with left flank tumours are infiltrated by higher numbers of neoantigen-specific CD4^+^ T cells (15). DCs are scarce and immature within solid tumors and are generally considered to exert their primary effects in lymph nodes (2, 7, 8, 16, 17). Because structural tissue cells greatly outnumber professional antigen presenting cells, express immune genes (18) and can be induced to present antigens (12, 19), we hypothesized that they are required for local antigen presentation and antitumor immunity.

Fibroblasts have been long considered as immunosuppressive cells (20). Thus, the recently identified MHCII antigen presenting cancer-associated fibroblasts (apCAFs) in pancreatic adenocarcinoma (PDAC) and breast carcinoma (BC) were presumed to induce immune tolerance and tumor escape (21–23). Here we show in murine models of lung cancer that fibroblast-specific targeted ablation of MHCII impairs local immunity, accelerating tumor growth. In primary human lung tumors CD4 T cells accumulate in apCAF dense spots. Human apCAFs directly primed adjacent effector CD4 T cell via the TCR and at the same time protected T cells from apoptosis via the c1q receptor c1qbp. These findings reposition apCAFs as immunostimulatory cells and propose a new functional node in the cancer-immunity cycle: CD4 T cells must receive a second wave of antigen presentation after their egression from draining lymph nodes within tumors for effective MHCII immunity to occur.

## RESULTS

### Antigen-presenting CAFs (apCAFs) prime adjacent CD4 T cells within human lung tumors

We undertook a pilot analysis of MHCII expression in human lung adenocarcinomas and squamous cell carcinomas by immunohistochemistry. We noticed abundant niches of elongated MHCII^+^ fibroblast-like cells inside the tumor bed **(Supplementary Figure 1)**. To elaborate on the fibroblastic identity of these MHCII^+^ cells we enzymatically dispersed primary human tumors and analyzed known mesenchymal markers by FACS **(Figure 1a)**. First, FACS impressively underestimated the frequencies of fibroblasts compared to imaging. This discrepancy likely reflected differences in detachment efficiency of different cell types following tissue digestion, with fibroblasts being more adherent to the extracellular matrix than immune cells and thus more difficult to disconnect. MHCII^+^ CAFs (apCAFs) were phenotypically similar to their MHCII^−^ counterparts and largely co-expressed FAP, PDGFRa and Podoplanin. Vimentin was expressed at variable levels, while aSMA was lowly expressed **(Figure 1a)**. 2D visualization in t-SNE plots (input all mesenchymal markers) confirmed that apCAFs do not cluster separately from MHCII^−^ CAFs. To determine the spatial relationship between apCAFs and CD4 T cells we analyzed whole tumor tissue sections by immunofluorescence staining. apCAFs were identified as cells that co-expressed MHCII and FAP. Quantitative analysis showed higher numbers of intratumoral MHCII^+^FAP^+^ relatively to MHCII^+^FAP^−^ cells, confirming the relative abundance of apCAFs among antigen presenting cells. T cells accumulated in regions with dense apCAFs and were identified closer to apCAFs rather than FAP^−^ MHCII^+^ cells **(Figure 1b)**. We also found a significant correlation between numbers of CD4 T cells and apCAFs in different regions of the same tumor section **(Figure 1b)**. This suggests that apCAFs create functional spots within lung tumors that sustain CD4 T cell populations and thus MHCII immunity.

**Figure 1.**
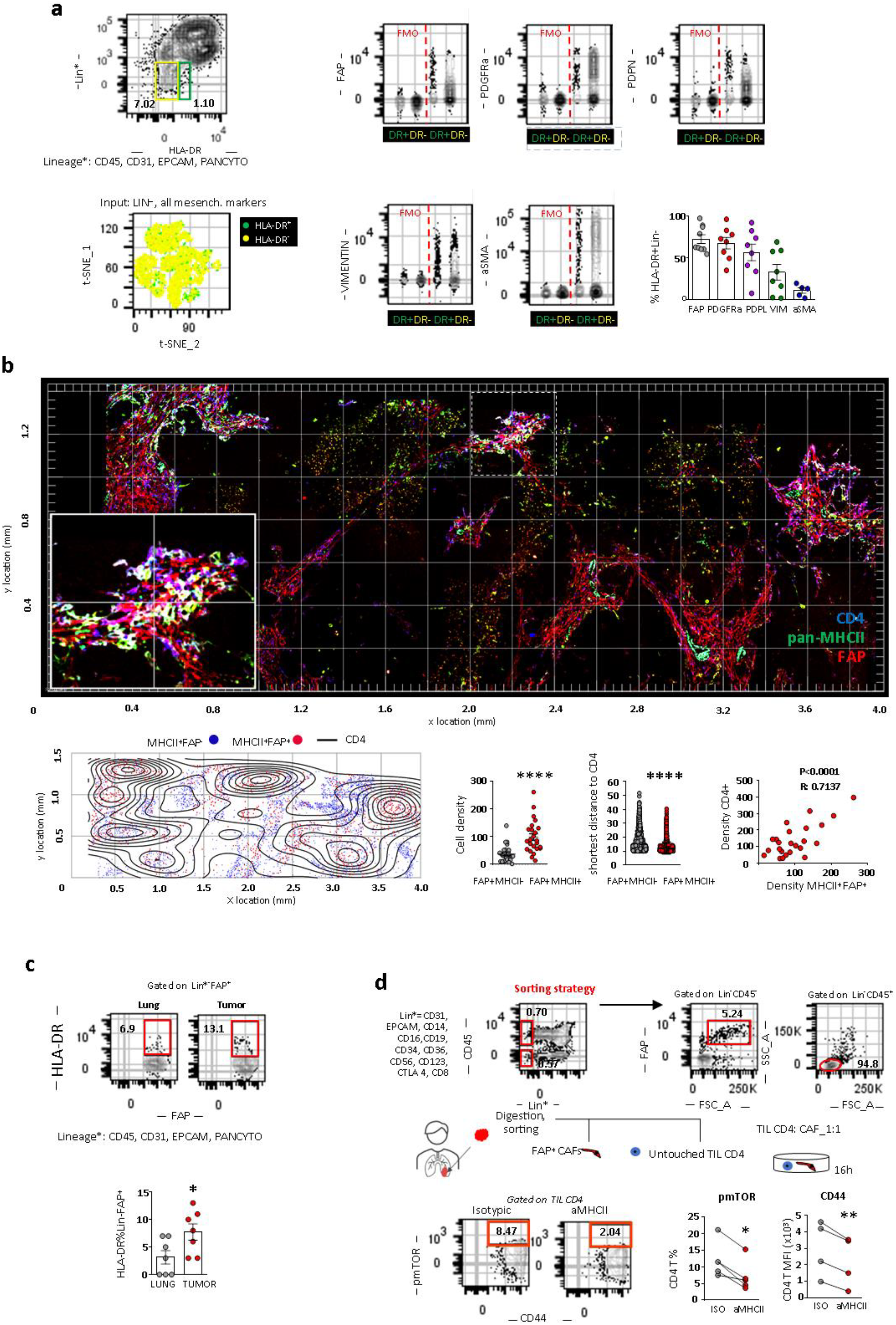
MHCII fibroblasts form CD4 T cell priming spots within human lung tumors. **a)** Representative FACS plots of a digested human lung tumor and expression of mesenchymal specific genes. Cumulative data (n=5-9 patients). **b)** Representative whole-slide imaging for panMHCII, FAP and CD4 and cellular spatial relationship map from a lung cancer patient. After acquiring XY coordinates, XY location of MHCII+FAP+ and MHCII+FAP-cells were overlaid with CD4 cell density contour. Distance between MHCII+FAP+ or MHCII+FAP- and the closest CD4 cell. Correlations between MHCII+FAP+ densities with CD4+ cell densities (cells per 25×10^4^μm^2^ regions in whole slide images, representative of n=3 patients). **c)** FACS plots showing MHCII in FS^hi^Lin^−^FAP^+^PDGFRa^+^ fibroblasts of paired digested human lung tumors and tumor-free lungs (n=7 patients). **d)** CD4 T cells and fibroblasts were sorted from the same tumor fragment and co-cultured with panMHCII blocking antibody or isotype control. Representative FACS plots and cumulative data on phospho-mTOR and CD44 (n=4 patients). **b, c, d)**\*P<0.05, **P<0.01, Unpaired or paired t-test.

We interrogated whether we could identify MHCII fibroblasts in healthy lungs. To rule out the possibility of contamination by non-fibroblastic cell populations we gated fibroblasts as FS^high^Lin^−^FAP^+^ cells. MHCII was expressed by a subset of normal lung fibroblasts, but it was more frequently detected in CAFs, suggesting that the tumor microenvironment drives and/or sustains apCAFs **(Figures 1c)**.

Spatial heterogeneity of T cell clones suggests regional activation by neighboring antigen-presenting cells that present their cognate antigens (13). We reasoned that apCAFs directly presented cancer MHCII peptides to adjacent CD4 T cells. To test this, we co-cultured primary human CD4 T cells and FAP^+^ CAFs, purified from the same lung tumour fragment and assessed phosphorylation of the key TCR/CD28 signaling node, mTOR **(Figure 1d)**. Primary cell phenotypes are dramatically altered once they are isolated and cultured. To avoid this bias, we used freshly FACS sorted primary tumor fibroblasts and freshly sorted autologous primary tumor infiltrating T cells for all our co-cultures. We used the bulk of CAFs rather than MHCII^+^ CAFs to avoid blocking TCR recognition and did not supplement the culture medium with exogenous cytokines. Approximately 1 in 10 intratumoral T cells responded to direct CAF contact by mTOR phosphorylation and CD44 up-regulation and these effects were abolished with MHCII blocking **(Figure 1d)**. Thus, human apCAFs acquire exogenous peptides within tumors in vivo and can directly prime adjacent CD4 T cells ex vivo.

### apCAFs are detected in tumors of IFNγ sufficient mice

To assess whether apCAFs were present in murine tissues we analyzed three models of orthotopic lung cancer. We inoculated one commercially available (LLC^mCherry^) or an autochthonous cancer cell line (CULA^zsGreen^) in the left lung lobe of syngeneic mice or injected melanoma cells (B16F10^mCherry^) in the tail vein of mice. Tumors were digested and subjected to FACS analysis. Intratumoral FS^high^CD45^−^EPCAM^−^ CD31^−^IAb(MHCII)^+^ non-cancer cells co-expressed podoplanin and PDGFRa, but not aSMA **(Figure 2a)**. Murine FAP antibodies showed nonspecific and unreliable staining and were thus excluded from the analysis. Similar to human apCAFs, 2D visualization in t-SNE plots indicated that apCAFs are admixed with MHCII^−^ CAFs. Immunofluorescence staining detected MHCII^+^Podoplanin^+^ cells within the tumor bed **(Figure 2a)**. Similar to what we had observed in humans, MHCII was more frequently expressed by CAFs relatively to healthy lung fibroblasts **(Figure 2b).** To determine whether fibroblasts depended on IFNγ for MHCII expression we inoculated LLC cells in the lungs of IL12 p35−/−, IL12p40−/−, IFNγR−/− and IFNγ−/− mice. MHCII expression was decreased in CAFs of IL12 deficient mice and severely impaired in those of IFNγ/IFNγR deficient versus wild type mice **(Figure 2b)**.

**Figure 2:**
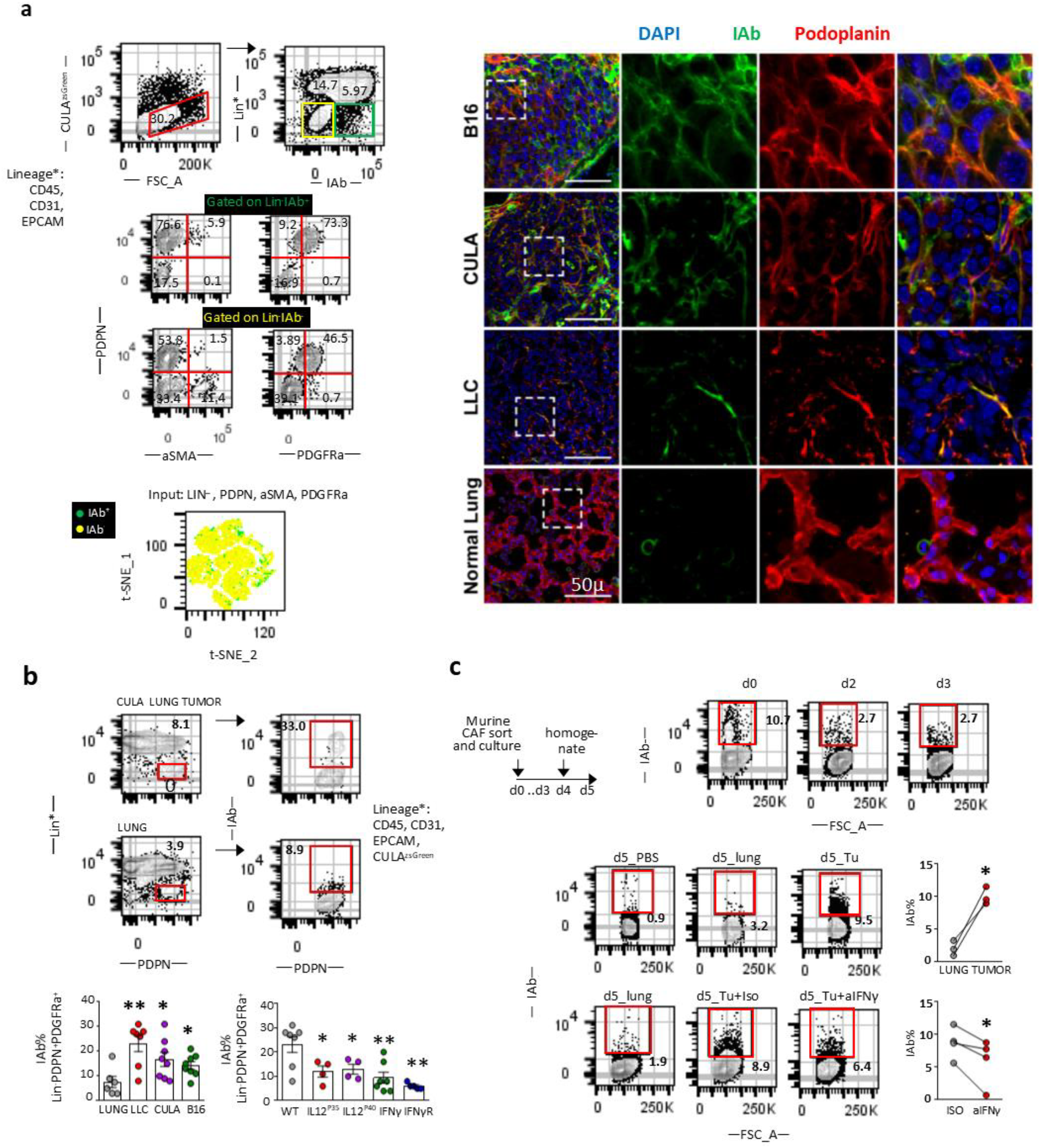
MHCII expression in fibroblasts is IFNγ-dependent. LLC/CULA lung adenocarcinoma cells or B16F10 melanoma cells were injected in the left lung lobe of syngeneic mice or through the tail vein, respectively. **a)** Representative FACS plots of a digested mouse CULA lung tumor and expression of mesenchymal specific genes. Immunofluorescence (IF) for IAb (MHCII) and podoplanin identifies MHCII fibroblasts. **b)** Up. Representative FACS plots of IAb in FS^hi^Lin^−^Podoplanin^+^PDGFRa^+^ fibroblasts of a CULA lung tumor and a healthy lung of wild type Bl6 mice. Down, left. Cumulative data on IAb expression of LLC, CULA and B16 lung tumors and healthy lungs of wild type Bl6 mice (n=6-8 per group). Down, right. Cumulative data on IAb expression of LLC lung tumors of cytokine-deficient mice (n=4-7 per group). **c)** FS^hi^Lin^−^Podoplanin^+^PDGFRa^+^ fibroblasts were isolated from pooled LLC lung tumors and cultured in 3D, as indicated. (n=4 experiments). **b, c)** *P<0.05, **P<0.01, Unpaired or paired t-test.

To probe the contribution of the tumor microenvironment in sustaining MHCII in fibroblasts, we developed an in vitro model system where we sorted PDGFRa^+^Podoplanin^+^ fibroblasts from LLC^mCherry^ tumors and cultured them in vitro. Although by day 3 all cultured CAFs had lost MHCII, exposure to fresh lung tumor, but not to healthy lung homogenates, restored MHCII expression **(Figure 2c)**. It should be noted that MHCII induction was observed upon 3D, but not 2D culture conditions, consistent with previous observations that 2D cultures induce MHCII loss in PDAC fibroblasts and differentiation to myofibroblasts (21). Notably, neutralizing IFNγ in tumor homogenates restrained MHCII induction in primary CAFs **(Figure 2c)**.

### Human and murine apCAF transcriptomes reveal conserved immune phenotypes and epithelial signature

To characterize human antigen-presenting fibroblasts we leveraged a public single-cell RNA sequencing (scRNAseq) dataset of primary human lung tumor, adjacent unaffected lung and computational analyses (24). There is no standardized computational method for characterizing gene expression levels in a binary form of “high” and “low”. Furthermore, gene expression levels correlate variably with corresponding protein levels. We hypothesized that we would be able to identify antigen presenting fibroblasts in single fibroblast RNA sequencing datasets if we used cDCs as a metric for physiologically relevant MHCII gene expression. Re-clustering of 1284 single-cells annotated as either fibroblasts (798) or cross-presenting cDCs (486) (E-MTAB-6149 and E-MTAB-6653, n=3 patients) via k-means clustering (input all MHCII genes, k=3 clusters supported by Silhouette scoring), separated fibroblasts into 2 clusters that shared (MHCII^+^) or did not share (MHCII^−^) MHCII expression features with cDCs. Similar to FACS analysis, dimensionality reduction (input 1785 expressed genes) showed that MHCII^+^ fibroblasts are predominantly admixed with MHCII^−^ fibroblasts and do not form a separate cluster **(Figure 3a)**. Differential expression analysis (DEA) revealed 115 deregulated genes in MHCII^+^ versus MHCII^−^ fibroblasts, 67 of which were upregulated **(Figure 3b)**. Among the top upregulated were signature genes of PDAC apCAFs (CD74, SLPI)(21), the IL6 gene (known as a prototype inflammatory CAF gene) (20), components of the complement pathway (CFD, C1QA, C1QB). Among the top down regulated genes was ACTA2 (encoding the myofibroblastic CAF gene aSMA) (20), a number of collagen genes (COL3A1, COL1A1, COL1A2) and other secreted extracellular matrix (ECM) proteins (SPARC, POSTN, BGN). Comparing pathway activities between MHCII^+^ and MHCII^−^ fibroblasts revealed enrichment in upregulated genes that are involved in immunological processes and down-regulated genes involved in ECM organization **(Figure 3b)**. Although MHCII^+^ fibroblasts were interspersed among MHCII^−^ fibroblast in t-SNE plots, it was striking that SFTC (encoding for surfactant protein C) was on the top of the list of up-regulated genes, raising the possibility that MHCII^+^ fibroblasts arise from alveolar type II cells, which constitutively express SFTC and MHCII (25). Three other genes known to be expressed by epithelial cells, CLU (clusterin) and the antimicrobial proteins lysozyme (LYZ) and SLPI, were amongst the 11 top up-regulated genes. Altogether our computational analysis of the transcriptome of MHCII^+^ fibroblasts depicts i) prominent immune functions, ii) a profile closer to inflammatory rather than myofibroblastic CAFs (IL6 high, a-SMA low, ECM proteins low) and iii) strong cues of an alveolar epithelial origin.

**Figure 3:**
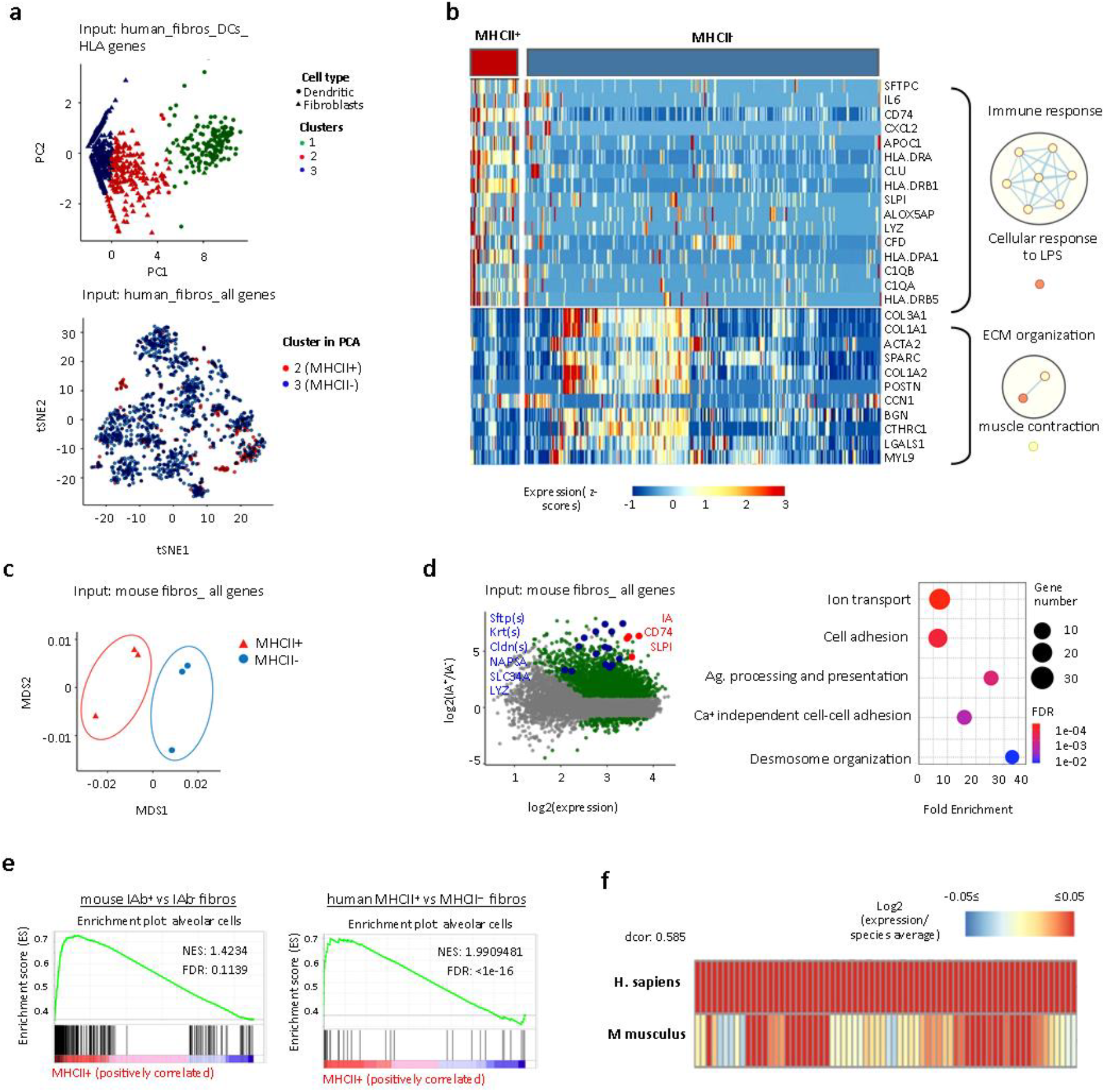
MHCII fibroblasts closely relate to alveolar epithelial across mice and humans. **a)** Fibroblast and dendritic cell clusters in scRNAseq datasets (E-MTAB-6149, −6653) from lung tumors and adjacent lungs were re-clustered to identify MHCII fibroblasts (n=3 patients). **b)** Heat map of differentially expressed genes (adj.P-values<0.05) between MHCII+ and MHCII-fibroblasts. GO enrichment analysis was performed using DAVID. Network visualization was conducted by Cytoscape’s EnrichmentMap. Nodes represent GO biological processes and edges connect functional terms with overlapping genes. Clusters of related nodes are circled and given labels reflecting broad biological processes. GO terms with an FDR < 0.1 are shown**. c)** MDS representation of bulk RNAseq data of sorted MHCII^+^ and MHCII^−^ fibroblasts from pooled mouse LLC lung tumors (n=3 experiments). **d)** Differential expression between MHCII^+^ and MHCII^−^ fibroblasts. Deregulated genes with adj.P-values<0.05 are indicated. GO enrichment analysis using upregulated genes with |Log2FC|>3 as input. **e)** GSEA enrichment plots for alveolar epithelial gene sets. **f)** Homology analysis of gene expression across mouse and human MHCII fibroblasts.

To investigate whether the human lung MCHII fibroblast signature was conserved across mice, we sorted MHCII^+^ and MHCII^−^ CAFs and performed bulk RNA sequencing **(Figure 3c**). Differential expression analysis (input all expressed 15675 genes, adj.P<0.05, |Log2FC|>3) revealed a large number of de-regulated genes (486), almost all of which (483) were up-regulated in MHCII^+^ CAFs **(Figure 3d**). As in humans, among the top upregulated genes were the 2 MHCII CAF signature genes CD74 and SLPI. Pathway analysis was in concordance with the immune-related pathways that were identified in our human dataset **(Figure 3d).** Intriguingly, the alveolar type II specific genes SFTPs and many alveolar epithelial cell genes were again found increased (CLAUDINs, KRTs, SLC34A, NAPSA, LYZ). Alveolar type II cells are long-lived cells that self-renew locally (26). To further support the hypothesis that MHCII fibroblasts originate from alveolar epithelial cells, we used alveolar cell type signature gene sets (curated from cluster markers, Supplementary Methods) and used them as inputs for GSEA. GSEA comparing MHCII^+^ versus MHCII^−^ fibroblasts of mice and humans demonstrated that MHCII^+^ fibroblasts were significantly enriched in genes that characterized alveolar epithelial cells in both species **(Figure 3e)**. To align the gene signatures expressed across MHCII fibroblasts in mice and humans, we coclustered genes based on their transcriptional profile in each species, using genes that were conserved and variable across MHCII fibroblasts in the human dataset **(Figure 3f)**. Our analysis denoted conservation of the MHCII fibroblast profile across the two species. Altogether these results indicate that the process of epithelial-to-mesenchymal transition regulates the in situ differentiation of alveolar type II cells to a subset of fibroblasts with antigen presenting functions.

### Deletion of MHCII in fibroblasts has a localized impact on tumor infiltrating CD4 T cells

We asked whether cancer antigen presentation by CAFs was a bystander, suppressor or driver of tumor progression in vivo. We set up pilot experiments starting with three murine lines expressing Cre recombinase (Cre) driven off Twist(27), collagene VI(28), and collagen 1a2 (Col1a2) promoters(29), which are lineage markers for mesenchymal cells. Lineage-restricted expression in LLC lung tumors was determined by Cre-driven GFP expression. An important fraction of intratumoral Twist-driven GFP expressing cells were CD45^+^ cells consistent with cells of the hematopoietic lineage, while Collagene VI-driven GFP-expressing cells could not be identified in lung tumors (**Supplementary Figure 2A**). Col1a2-driven GFP-expressing cells in lung tumors of Col1a2-CreER mice were CD45^−^CD31^−^EPCAM^−^ and largely Podoplanin^+^ -indicating that they were bona fide fibroblasts (**Supplementary Figure 2B**). Therefore, we used Col1a2-CreER mice for our subsequent studies. IA^b^ is the MHC class II molecule expressed in B6 mice and its expression can be deleted from Cre-expressing cell lineages using a floxed IAb gene (I-A^b-fl/fl^) (30). We inoculated LLC^mCherry^ cells that expressed the model antigen ovalbumin (ova) in the lungs of Col1a2 CreER^+^IAb^fl/fl^ and Col1a2 CreER^−^IAb^fl/fl^ mice. Lung tumor burden was increased and tumors appeared immunologically colder, with fewer CD4 T cell, OVA-specific CD8 T cell and B cells (**Figure 4a**). Importantly, CD4 T cell-depleting antibodies abolished differences in tumor burden between Col1a2 CreER^+^IAb^fl/fl^ and Col1a2 CreER^+^IAb^fl/fl^ mice, indicating that MHCII fibroblasts were strongly dependent on CD4 T cells for anti-tumor function **(Figure 4b)**. RNAseq analysis of FACS sorted TIL CD4 showed a high number of down regulated genes in Col1a2 CreER^+^IAb^fl/fl^ versus Col1a2 CreER^−^ IAb^fl/fl^ mice **(Figure 4c)**. Among top downregulated were genes involved in gene transcription (CRCT2, KAT7, EP300), cellular energy metabolism (GLUL, LIPA, PRKAG2), protein ubiquitination and proteosomal degradation (PSMD6, CDC26, TRIM26) (**Figure 4c**). Signature genes of Th1, Th2, Th17, Th9 and Tregs, cytotoxicity or exhaustion marker genes were not different between the two groups **(Supplementary Figure 3)**. Likewise, pathway activities of downregulated genes showed significant enrichment in macromolecular metabolic processes. TCR activation is known to play a fundamental role in T cell metabolism (31). Taken together, the results described above suggest that antigen presentation by MHCII fibroblasts is a critical driver of MHCII immunity in vivo.

**Figure 4:**
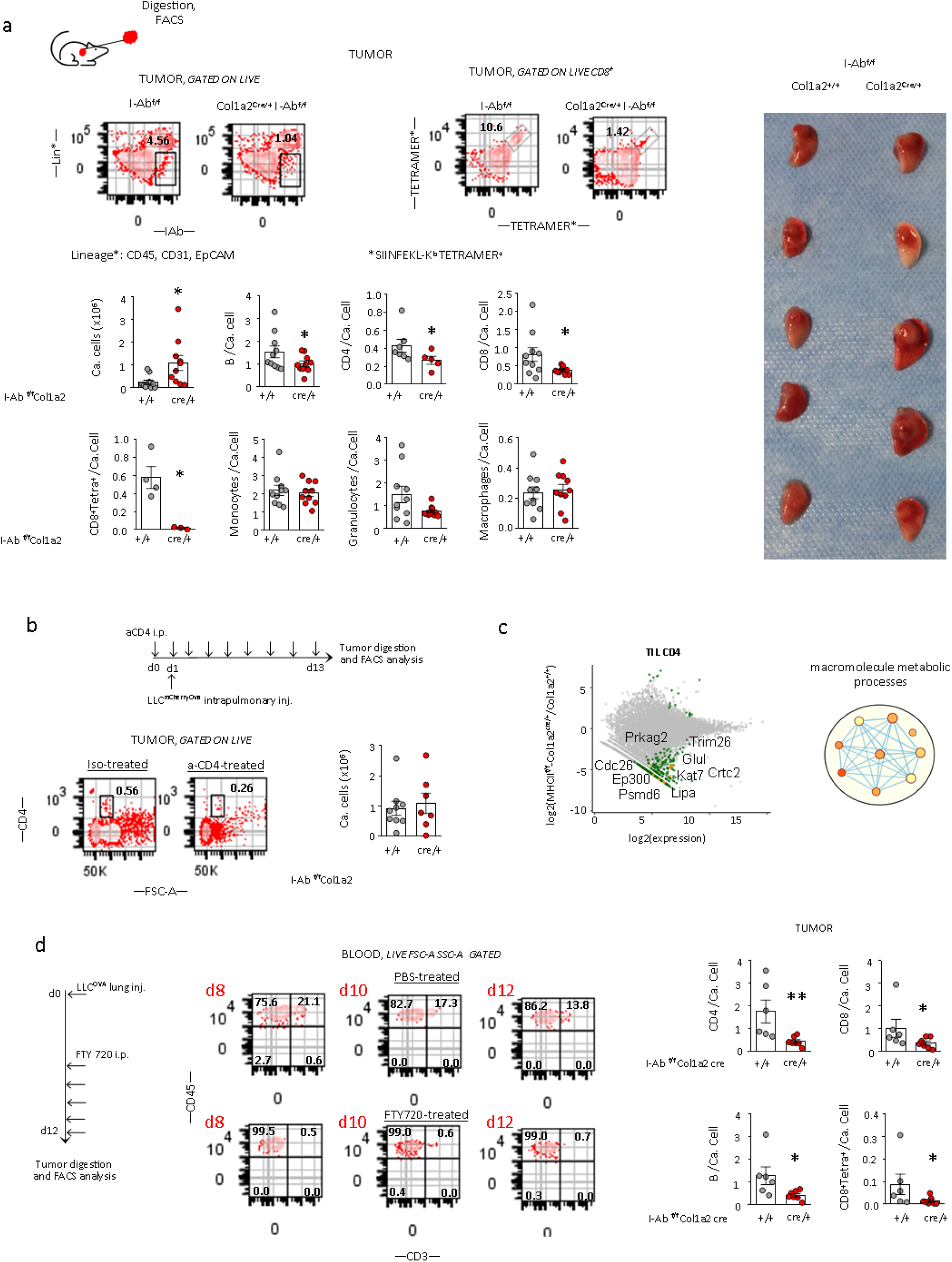
Local antitumor immunity depends on MHCII-peptide presentation by CAFs. **a)** Up. Col1a2 CreER^+^I-Ab^fl/fl^ and Col1a2 CreER^−^I-Ab^fl/fl^ mice were inoculated with LLC^mcherryOva^. Representative FACS plots of Lin^−^IAb^+^ and CD8^+^tetramer^+^ cells in digested tumors. Down. Cumulative data of tumor burden and intratumoral immune profiles (n=3-7 per group, pooled from 2 experiments). Right. Representative photograph. **b)** As in A, under CD4 T cell depleting antibody or isotype control treatment (n=3-6 per group, pooled from 2 experiments). **c)** Differential expression between purified intratumoral CD4 T cells from Col1a2 CreER^+^I-Ab^fl/fl^ versus Col1a2 CreER^−^I-Ab^fl/fl^ mice (n=7). Deregulated genes with FDR<0.01 are indicated. GO enrichment analysis was performed using DAVID. Network visualization was conducted by Cytoscape’s EnrichmentMap. Nodes represent GO biological processes and edges connect functional terms with overlapping genes. Clusters of related nodes are circled and given labels reflecting broad biological processes. GO terms with an FDR < 0.1 are shown. **d)** As in A, under FTY720 treatment (n=6-7 per group, pooled from 2 experiments). **a, b, d)** Mann-Whitney U test. P-value<0.05

Lymph node fibroblastic reticular cells (FRCs) have contradictory roles in immune responses, such as inhibition of T cell proliferation via nitric oxide (32) and T cell reprogramming towards memory T cells via IL6 (33). Albeit FRCs in LNs acquire transcriptional programs typically associated with tumor escape (34), we considered the possibility that immune evasion in Col1a2 Cre^+^IAb^fl/fl^ mice resulted from MHCII deletion in FRCs. We assessed MHCII expression in LN FRCs of Col1a2 CreER^+^IAb^fl/fl^ versus Col1a2 CreER^−^IAb^fl/fl^ mice and found a trend toward decreased MHCII^+^ FRCs **(Supplementary Figure 4)**. However, there were no gross differences in nodal CD4^+^, CD8^+^ T cells and B cells **(Supplementary Figure 4)**. We questioned whether antigen presenting CAFs could sustain MHCI and MHCII immunity in the absence of continuous T cell migration from LNs. Therefore, we treated mice with a S1PR antagonist FTY720, which blocks egress of T cells from LNs, for 5 consecutive days, starting on day 7 after OVA-LLC transplantation^30^. CD4 and CD8 T cells were again found decreased, while ova peptide-MHCI tetramers stained less of CD8 T cells in lung OVA-LLC tumors of FTY720-treated MHCII conditional knockout mice **(Figure 4d)**. These results suggest that targeting MHCII in fibroblasts facilitates immune escape by acting locally and that T cell immunity in Col1a2 Cre^+^IAb^fl/fl^ mice is compromised by MHCII loss in CAFs rather than FRCs.

### apCAFs rescue intratumoral CD4 cells from apoptosis via the c1q receptor c1qbp

Effector T cells have less stringent and incompletely understood activation requirements compared to naive T cells(35). Since we could barely detect any of the classical co-stimulatory molecules (CD80, CD86, CD40) in MHCII fibroblasts we mined their single cell transcriptome looking for other immune-related molecules with ligand-receptor functions. Human MHCII^+^ fibroblasts overexpressed TYROPB and CD47 which ligate to SIRP receptors, CD52 and ANXA1 which bind to SIGLEC10 and ALX/FPR2, respectively, and C1q which binds to cC1qR/calreticulin and gC1qR/C1qbp (**Figure 1I**). None, but the receptor of the globular head of C1q, i.e. gC1qr/C1qbp, was found to be expressed on the surface of primary human TIL CD4 cells **(Figure 5a)**. C1qbp is located in most cell types in the mitochondria where it is responsible to maintain transcription of mitochondrial proteins. In the cytoplasm it acts as a regulator of RNA stability and plays a critical role in mRNA splicing. The role of c1q as an extracellular signaling molecule and c1qbp as a plasma membrane receptor on T cells has been rarely reported (36–38) and is largely unexplored. We set out to investigate whether MHCII fibroblast-derived c1q amplified TCR signaling on CD4 T cells via membrane c1qbp. First, we confirmed increased c1q gene expression by MHCII^+^ versus MHCII^−^ lung tumor fibroblasts in distinct patients **(Figure 5a)**. Then, we co-cultured human effector CD4 TILs and CAFs in the presence of C1qbp blocking antibodies. Surprisingly, we did not observe any effect on TCR signaling, but a decrease in the numbers of viable T cells (**Figure 5b**). This lead us hypothesize that c1q is an extrinsic pro-survival signal for CD4 T cells. To test this, we cultured stimulated peripheral blood CD4 T cells for 1 h under serum starvation conditions. Purified c1q acted directly on T cells to almost completely rescue them from apoptosis **(Figure 5c)**. Depleting MHCII fibroblasts from the CAF-TIL CD4 co-cultures abrogated the apoptotic effect of aC1qbp, indicating that ac1qbp blocks MHCII fibroblast-derived c1q **(Figure 5d)**. We consider unlikely that these results have been confounded by an autocrine effect of T cell intrinsic c1q because we found lower c1q gene expression levels in TIL CD4 than MHCII^+^ CAFs and aCD3/aCD28 stimulation did not upregulate CD4 T cell-intrinsic c1q **(Supplementary Figure 5)**. Based on these findings, we examined whether c1qbp could function as pro-survival receptor in vivo. OVA-specific OTII T cells were transduced with a c1qbp versus empty lentivirus and adoptively transferred in LLC-OVA lung tumor bearing mice. An increased number of C1qbp-OTII cells were identified 3d post adoptive transfer within lung tumors and this was accompanied by a decrease in tumor burden **(Figure 5e)**. Altogether these data suggest that CD4 T cells depend on c1qbp for their survival within tumors and point to MHCII fibroblasts as a potent c1q source.

**Figure 5:**
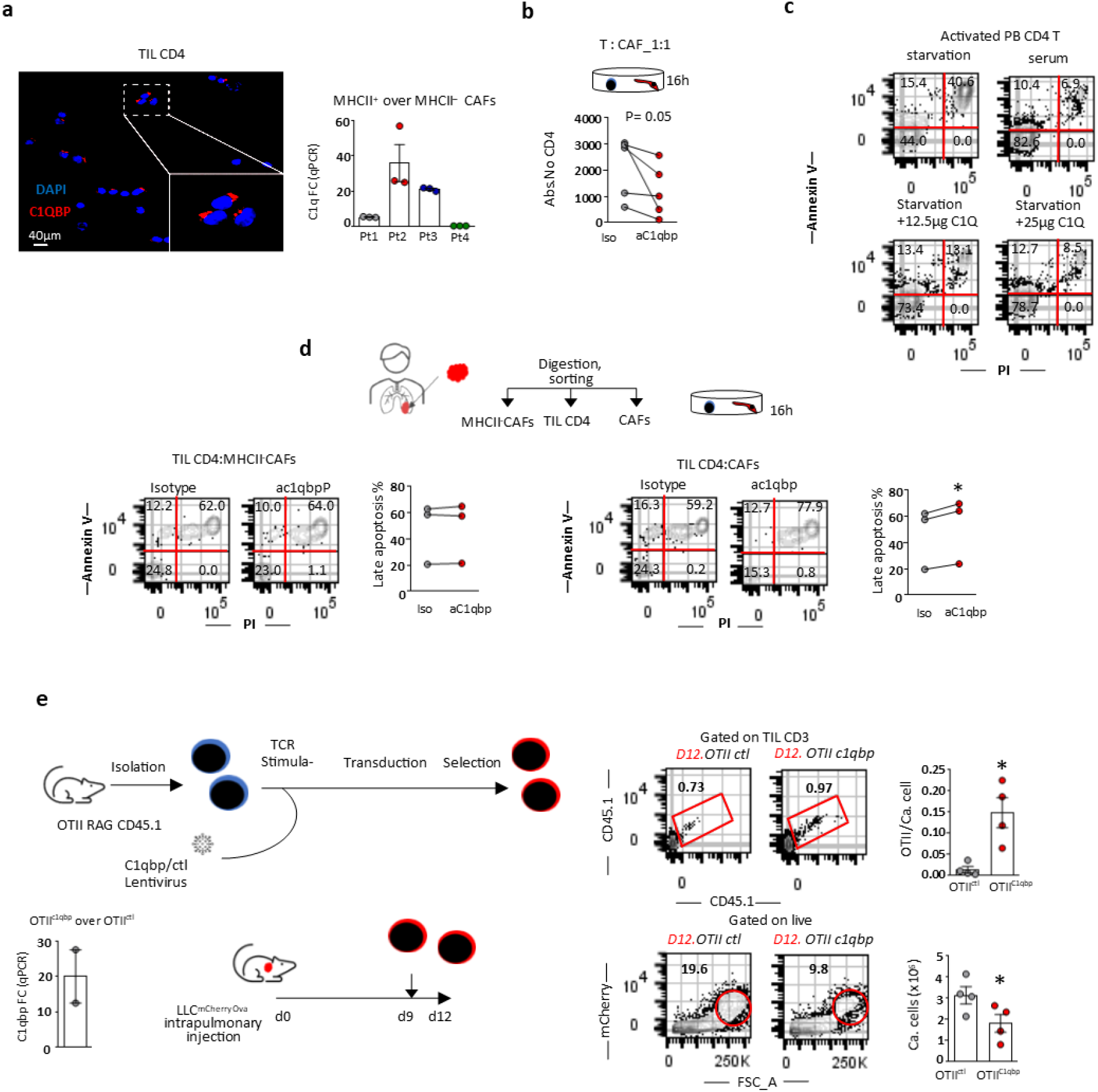
apCAF-derived c1q helps tumor infiltrating CD4 T cells evade apoptosis. **a)** Immunofluorescence (IF) for c1qbp shows surface expression in intratumoral CD4 T cells. qPCR for c1qb in sorted MHCII^+^ versus MHCII^−^ FS^hi^Lin^−^FAP^+^ CAFs in human lung tumors (n=4 patients). **b)** CD4 T cells and CAFs were sorted from the same tumor fragment and co-cultured with ac1qbp blocking antibody or isotype control. Abs .numbers of live T cells were assessed by FACS (n=5 patients). **c)** Peripheral blood (PB) CD4 T cells from a healthy donor were sorted and activated with aCD3/aCD28 beads prior to undergoing serum starvation with or w/o purified c1q. Representative FACS plots of 2 experiments. **d)** CD4 T cells, total CAFs and MHCII^−^ CAFs were sorted from the same tumor fragment. T-CAFs were co-cultured with ac1qbp blocking antibody or isotype control. T cell apoptosis was assessed using annexin-V and PI staining. Representative FACS plots and cumulative data are shown (n=3 patients). **e)** C1qbp overexpressing OTII T cells were adoptively transferred in LLC^mcherryOva^ lung tumor bearing mice. Representative FACS plots of LLC^mcherryOva^ cells and OTII cells in digested lung tumors. Cumulative data (n=4 per group, pooled from 2 experiments). **a-e)** Unpaired or paired t-test. P-value<0.05

## DISCUSSION

A functional dichotomy is currently thought to exist between draining lymph nodes and tumors, whereby T cells are primed in lymph nodes and exert effector functions in tumors. Here we show that antitumor effector T cells require secondary in situ TCR stimulation by a subset of CAFs for effective immunity to occur. Single cell/bulk RNA sequencing approaches revealed a conserved apCAF transcriptome signature among lung tumors of mice and humans that aligned to that of PDAC apCAFs and was enriched for immune-related and alveolar epithelial genes, suggesting that apCAFs arise from epithelial cells through the process of epithelial-to-mesenchymal transition. Fibroblast-specific targeted ablation of MHCII induced a lymph node-independent impairment in local immunity, accelerating tumor growth. In primary human lung tumors CD4 T cells preferentially homed dense MHCII fibroblast areas. Primary human fibroblasts directly activated the TCRs of tumor infiltrating CD4 T cells and at the same time protected them from apoptosis via the c1q receptor c1qbp, which unveiled a previously unrecognized function of c1q/c1qbp as an anti-apoptotic ligand/receptor pair for CD4 T cells. Our studies introduce non-hematopoietic cells as key cancer antigen-presenting cells and suggest that the final outcome of initial CD4 T cell priming within tumor draining lymph nodes depends on de novo antigen presentation within tumors.

In MHCII IHC tumor slides MHCII fibroblastic cells were not identified within macroscopically visible tertiary lymphoid structures (TLS) structures, suggesting that intratumoral fibroblastic antigen presenting niches are distinct to TLS. It is likely that that TLS are mainly sites of B cell somatic hypermutation and plasma cell generation (39–41), while intratumoral antigen presenting fibroblasts form immunological niches for T effector cells. MHCII fibroblasts did not express classical costimulatory molecules but were still able to activate the TCR/CD28 pathway in effector T cells and enhance T cell metabolism. We consider two possibilities that may explain this contradiction: i) Costimulatory and co-inhibitory receptors of effector T cells display great diversity and promiscuity. Thus MHCII fibroblasts mediate effector T cell activation via nonclassical costimulatory molecules (35). ii) Antigen recognition suffices to activate effector T cells (42).

Our results provide the first, to the best of our knowledge, demonstration that a complement component is an antiapoptotic factor. C1q is known to regulate basic metabolic processes of T cells (36–38) but has been disregarded as an extracellular signal. This is also related to the fact that the receptor of its globular head, C1qbp, has a very short intra-cytoplasmic tail and must complex with other membrane molecules to trigger intracellular signaling (43, 44). The outcome of T cell therapy is highly dependent on T cell survival within tumors (45, 46). We showed that C1qbp overexpression increases the numbers of adoptively transferred tumor specific CD4 T cells within tumors. Conceivably, it is likely that the increased efficacy of C1qbp-OTII cells is not only the result of apCAF-derived c1q. Nonetheless our results showing that i) apCAFs express high levels of c1q; ii) they are a key antigen presenting cells within lung tumors iii) c1q acts directly on T cells to rescue them from apoptosis; iv) TIL CD4 cells reside within apCAF spots, altogether suggest an important role for apCAF-derived c1q as a pro-survival factor for TIL CD4 cells.

The following arguments indicate that lung apCAFs derive from alveolar epithelial type II (ATII): i) in both human and mouse datasets the most up-regulated genes were surfactant proteins, which are highly specific for ATII cells, ii) constitutive MHCII expression is a hallmark of ATII cells(25), iii) ATII cells can transit to fibroblasts though the process of epithelial-to-mesenchymal transition (47, 48) iv) PDAC apCAFs also express epithelial specific genes (21). Lung apCAFs should be considered bona fide fibroblasts rather than ATII cells because of their characteristic fibroblastic morphology, their ability to easily grow in culture (primary alveolar epithelial cells are notoriously difficult to grow), the absence of surface EPCAM, the co-expression of multiple signature mesenchymal genes (49). We have rejected the possibility that apCAFs originate from cancer cells. In the human dataset tumor cells clustered away from fibroblasts, while in the mouse data set cancer cells were brightly fluorescent and negatively selected. Furthermore, we never detected fluorescence in murine fibroblasts that were cultured up to passage 5.

The apCAFs that were recently identified in PDAC and BC were presumed to induce T cell anergy or differentiation to T regulatory cells (21–23). This presumption opposed the increase in T regulatory cells that had been observed upon fibroblast deletion in PDAC (50). In our models MHCII deletion in fibroblasts did not alter CD4 T cell subsets. However, it caused a profound decrease in the metabolic rate of tumor infiltrating CD4 T cells. This finding fits perfectly well with the pivotal role of T cell metabolism in antigen-specific responses (31). Further supporting the idea that non-hematopoietic cells can act as stimulating rather than tolerizing antigen presenting cells, MHCII intestinal epithelial cells activate CD4 T cells in graft-versus-host disease and diverse types of non-hematopoietic lung antigen-presenting cells prime T resident memory (TRM) cells (12, 19). IFNγ drives MHCII expression in epithelial cells (19). Our results also demonstrate that the tumor microenvironment controls MHCII in fibroblasts via IFNγ. That said, positive clinical responses to immune checkpoint blockade are associated with increased IFNγ and MHCII expression (51–53). Thus, on the basis of our models, another mechanism behind the success of checkpoint inhibitors could be their ability to sustain IFNγ-dependent apCAF spots which in turn activate the unleashed effector T cells within tumors.

Current immunotherapeutic strategies largely focus on immune cells. Our works point to mesenchymal cells as key partners in cancer immunotherapies. We envisage two possible scenarios for the immunotherapeutic exploitation of our findings: in vivo delivery of tumor antigens to apCAFs and amplification of the antigen presenting fibroblastic spots by cellular re-programming of resident epithelial cells to transition to fibroblasts in situ. Conceivably, the above strategies should be combined with IFNγ-inducing therapies, such as checkpoint inhibitors, that will sustain MHCII expression within tumors.

## Acknowledgments

We would like to thank G. Kollias for critical insights and discussions. G. Kollias and P. Veginis for reviewing the manuscript. We acknowledge G. Kollias, V. Koliaraki, E. Andreakos, J.P. Gorvel, and G. Stathopoulos for sharing mouse models, cell lines and protocols. The NIH Tetramer Core Facility for SIINFEKL tetramers. We thank BSRC “Al. Fleming” flow cytometry, animal house, imaging and genomics facilities and the bioinformatics facility of “The Greek Research Infrastructure for Personalised Medicine (pMedGR)” (MIS 5002802). The study was supported by a Hellenic Foundation for Research and Innovation Grant (HFRI-1289) and a “Fondation Santé” research grant to MT. DK is supported the Greek State Scholarships Foundation (MIS-5033021).

## Author contributions

Conception and design study, M.T.; Development of methodology, M.T.; Acquisition of data, D.K., E.A., K.G., P.S. and A.P.; Analysis and interpretation of data, D.K., E.A., K.M.V., C.T., A.P. and M.T.; Providing human samples, I.V., E.K., K.V. E.S., K.P. and A.K.; Writing the manuscript, M.T.; Preparing the figures, D.K., E.A., K.M.V., K.V., P.S., C.T., A.P., M.T.; Study supervision, M.T.

## Competing Interests statement

None

## METHODS

### Human samples

Paired human lung tumour and macroscopically healthy lung resection specimens were obtained from patients with lung adenocarcinoma (LUAD) and squamous cell carcinoma (LUSC) at Sotiria Chest Hospital. The study was approved by the ethical committee of the Hospital and informed consent was obtained from all patients. LUADs and LUSCs were clinically scored and staged according to the International Union against Cancer (UICC) TNM staging system. The clinical and pathological characteristics of all patients included in this study are summarized (Supplementary Table 1).

### Mice

Col1a2-cre/ERT,-ALPP mice (Jackson Laboratories, ID #029235), Tg (Col6a1-cre)1Gkl (ColVI-Cre) mice (MGI:3775430), B6.129X1-Twist2tm1.1(cre)Dor/J (Twist2-Cre) mice (Jackson Laboratories, ID # 008712), mTmG mice (Jackson Laboratories, ID #007576), IL12 p35 knockout (Jackson Laboratories, ID #002691), IL12 p40 knockout (Jackson Laboratories, ID # 002693) were provided by G. Kollias, (BSRC Al. Fleming), OTII mice (Jackson Laboratories, ID #004194) by V. Andreakos (BRFAA), IFNgR (Jackson Laboratories, ID #003288) and IFNg knockout mice (Jackson Laboratories, ID #002287) from J.P. Gorvel (CIML). B6.129X1-H2-Ab1tm1Koni/J (I-AB-flox, ID #013181) and B6.SJL-Ptprc^a^ (CD45.1, ID #002014) mice were purchased from The Jackson Laboratories). For MHCII deletion in fibroblasts the Col1a2-cre mice were crossed to I-AB-flox mice that possess a loxP-flanked neo cassette upstream of exon 1, and a single loxP site downstream of exon 1 of the H2-Ab1 locus. For T cell responses against ovalbumin-expressing cancer cells, OTII mice that express the mouse alpha-chain and beta-chain T cell receptor that pairs with the CD4 co-receptor and is specific for the chicken ovalbumin 323-339 peptide in the context of I-Ab, were crossed to Rag1 knockout mice and mice that express the CD45.1 (Ly5.1 PTP) alloantigen. For reporter studies Col1a2-cre, ColVI-Cre and Twist2-Cre were crossed to mTmG mice.

### Cell lines

The Lewis Lung Carcinoma cell line (LLC), obtained from American Type Collection Cultures (Manassas, VA), the C57BL/6-derived urethane-induced lung adenocarcinoma CULA cell line, provided by G. Stathopoulos (Helmholtz Zentrum München) and the C57BL/6-derivered melanoma cell line B16F10, provided by V. Kostourou (BSRC Al. Fleming), were transduced with zsGreen and/or OVA^mCherry^ lentiviral vectors. Cells were maintained in DMEM, containing 10% heat-inactivated FBS, 1% L-glutamine and 1% penicillin/streptomycin (Gibco). All the cells were tested negative for the presence of mycoplasma contamination using a PCR-based technology. Cell lines were not authenticated.

### Sample preparation and staining

Human and murine tissue specimens were perfused with PBS and cut into pieces. Minced samples were immediately processed or cryopreserved using BioCool (SP Scientific) in Recovery Cell Culture Freezing Medium (Gibco, Cat. No.12648-010) enriched with 10% DMSO. Human and mouse tumour and lung tissue fragments were enzymatically digested in 10% FBS/HBSS (Gibco) using Collagenase IV (Sigma-Aldrich, 1 mg/ml, Cat. No.C7657), Dispase II (Roche, 1 mg/ml, Cat. No.SCM133) and Dnase I (Sigma-Aldrich, 0.09 mg/ml, Cat. No DN25) for 1 hour at 37°C with agitation. Murine lymph nodes were digested 10% FBS/HBSS using Collagenase P (Sigma-Aldrich, 1 mg/ml, Cat. No. 11 249 002 001), Dispase II (Roche, 1 mg/ml,) and Dnase I (Sigma-Aldrich, 0.09 mg/ml,) for 1 hour at 37°C with agitation. Human tumor cells were passed through a 100um cell strainer and murine tumor cells were passed through a 70um cell strainer. Murine splenocytes were mechanically dissociated by passing through a 40um cell strainer. Cells were washed with FACS buffer (2%FBS/PBS/1.5mM EDTA), centrifuged and re-suspended in FACS buffer. Non-specific binding was blocked by incubating cells with human anti-Fc Receptor antibodies (TrueStain, Biolegend, Cat. No. 101320) or anti-mouse CD16/32 Fc block (Biolegend, Cat. No. 101310).

Staining of human tissue was performed with the following antibodies (all from Biolegend, unless otherwise stated): CD45-PE, (Cat. No. 304007), CD31-PE, (Cat. No. 303105), EpCAM-PE, (Cat. No. 324205), FAP-Alexa700 (R&D Biosystems, Cat. No. MAB3715), Podoplanin APC-Fire 750, (Cat. No. 337023), CD140a (PDGFRa)-PE-Cy7, (Cat. No. 323507), CD152-PE (CTLA-4, Cat. No. 369603), CD40-BV421, (Cat. No. 334331), CD80-BV510, (Cat. No. 305233), CD86-APC, (BD,Cat. No. 555660), CD45-PE, (Cat. No. 304007), CD8-PERCP-Cy5.5, (Cat. No. 301031), HLA-DR-BV785 (Cat. No. 307641), HLA-DR-DP-DQ-APC (Cat. No. 361713), CD44-APC (Cat. No. 103011), CD14-PE (Cat. No. 301805), CD15-PE (Cat. No. 301905), CD16-PE (Cat. No. 302007), CD19-PE (Cat. No. 302207), CD34-PE (Cat. No. 343505), CD36-PE (Cat. No. 336205), CD56-PE (Cat. No. 304605), CD123-PE (Cat. No. 306005) and CD253-PE (Cat. No. 308215) for 30 minutes at 4°C.

Murine cells were stained with the following antibodies (all from Biolegend, unless otherwise stated): CD45-APC-CY7 (Cat. No. 103115), CD3-PE-CY7 (eBioscience, Cat. No. 25-0031-82), B220-PERCP (eBioscience, Cat. No. 45-0452-82), CD8-PERCP (Cat. No. 126610), CD4-FITC (BD Pharmigen, Cat. No. 553651), CD45-Alexa700 (Cat. No. 123128), CD31-Alexa700 (Cat. No. 102443), EpCAM-Alexa700 (Cat. No. 334331), CD140a (PDGFRa)-PE (Cat. No. 334331) or BV785 (Cat. No. 118239), Podoplanin-PE Cy7 (Cat. No. 127412), MHCII-APC-Cy7 (Cat. No.107627), CD45.1-Alexa700 (Cat. No. 110723), CD11b (BD Pharmigen, Cat. No.557397), CD11c (BD Pharmigen, Cat. No. 553802), CD19 (Cat. No. 115507), B220 (BD Pharmigen, Cat. No.553089), CD49b (Cat. No. 103506), CD105 (Cat. No. 110723), MHCII (Cat. No. 107607) and TER119 (Cat. No. 116207). Tetramers SIINFEKL-PE and SIINFEKL-APC were provided by NIH.

For intracellular staining, cells were fixed and permeabilised using the Intracellular Fixation & Permeabilization Buffer Set (eBioscience, Cat. No. 88-8823-88), followed by staining for human samples with PAN-Cytokeratin-PE (Sigma-Aldrich, Cat. No. SAB4700668), Vimentin-Alexa Fluor 674 (Abcam, Cat. No. ab194719), a-Sma-FITC (Sigma, Cat. No. F3777), and Alexa Fluor 647-anti-mouse (Invitrogen, Cat. No.A21235) secondary antibody. For murine samples, also a-Sma-FITC (Sigma, Cat. No. F3777) and Vimentin-Alexa Fluor 674 (Abcam, Cat. No. ab194719) were used.

For intracellular staining of phosphorylated proteins, cells were fixed and permeabilised using the Intracellular Fixation & Permeabilization Buffer Set (eBioscience, Cat. No. 88-8823-88) as per manufaturers’ recommendations for detection of intracellular phosphorylated proteins, followed by staining with pmTor-PE-Cy7 (eBiosciences, Cat. No. 25-9718-42).

Sytox-Green viability dye (ThermoScientific, Cat. No.S7020), Zombie NIR (Biolegend, Cat. No. 423105), Zombie Violet (Biolegend, Cat. No. 423113) or BD Horizon, Fixable, Viabilility, Stain 700 (BD Pharmigen, Cat. No. 564997) were used to exclude dead cells.

For Annexin V/ PI stain up to 5×10^4^ cells were resuspended in Binding Buffer (0.01M Hepes, pH 7.4, 0.14M NaCl, and 2.5 mM CaCl2) and stained with Annexin V FITC (Cat. No. 640906) for 30 minutes at 4°C. Cells were washed and resuspended in propidium iodide (PI)-Binding Buffer solution (0.1mg/ml, Sigma, Cat. No. P4170).

### Functional ex-vivo assays

For human immunological assays, CAFs and CD4^+^ T cells were harvested from the same tumour fragment and dispersed into single-cell suspensions as stated above. CAFs were sorted as FSC-A^High^CD45^−^CD31^−^EPCAM^−^FAP^+^PDGFRa^+^ cells. To preserve functional TCRs, intratumoral CD4^+^ T cells were sorted as FSC-A^Low^SSC-A^Low^CD45^+^CD14^−^ CD15^−^CD16^−^CD19^−^CD34^−^CD36^−^CD56^−^CD123^−^CD8^−^CTLA-4^−^ cells. CAFs were co-cultured overnight with CD4^+^ T cells at a 1:1 ratio in the presence of pan-HLA antibody (5μg/ml, Biolegend, Cat. No.361702) or GC1q R antibody (10μg/ml, Abcam, Cat. No. ab24733) or isotypic controls (mouse IgG2a (Biolegend, Cat. No.361702) and mouse IgG1 (Abcam, Cat. No. ab170190) respectively). Cells were co-cultured in complete RPMI (Gibco) medium.

Classical Ficoll-Pague (StemCell, Cat. No. 07861) density gradient centrifugation was followed for PBMCs isolation from human peripheral blood. Untouched CD4^+^ T cells were negatively selected from PBMCs using anti-PE MicroBeads (Miltenyi Biotech) after incubation with the following PE-conjugated antibodies: CD14, CD15, CD16, CD19, CD34, CD36, CD56, CD123, CD253 and CD8 according to manufacturer’s instructions. Human T Cell Activation/Expansion Kit (Miltenyi Biotech) was utilized for activation of CD4^+^ T cells according to manufacturer’s protocol. T cells were cultured in Click’s medium supplemented with 10% heat-inactivated human serum (Sigma-Aldrich, H4522), 1% L-glutamine and 1% penicillin/streptomycin, 1% sodium pyruvate, 1% MEM non-essential amino acids, 2mM HEPES, β-mercaptoethanol and 50U/ml huIL-2 (Peprotech, Cat. No. 200-02). For selected experiments T cells were cultured in serum-depleted medium with or without native human C1q protein (100ug/ml, Abcam, ab96363).

For murine 3D CAF cultures, CAFs sorted as FSC-A^High^CD45^−^CD31^−^EPCAM^−^PDPN^+^PDGFRa^+^ cells were resuspended in liquified Matrigel (354254, Corning) at 4°C. Approximately 30.000cells/well were seeded in a 25ul drop of Matrigel in the center of wells of a 48-well tissue culture plate. The matrix was allowed to set for 15 minutes at 37°C before adding 400ul complete RPMI into each well. At day 5 of culture, RPMI medium was replaced with fresh RPMI supplemented with 30% healthy lung or tumor lung homogenate plus/minus anti-IFN-γ (10ug/ml, Biolegend, Cat. No. 505705). To obtain tissue homogenates, tumors or healthy lungs were excised, weighed and homogenized in PBS (30% weight/volume) on ice. Samples were centrifuged at 13000rpm for 10 min at 4oC and supernatants were added in wells containing matrigel encapsulated cells.

### In vivo studies

Gender and age-matched, over 8-week-old mice were used for all studies. All mice were housed under standard special pathogen-free conditions at BSRC Alexander Fleming. All animal procedures were approved by the Veterinary Administration Bureau, Prefecture of Athens, Greece under compliance to the national law and the EU Directives and performed in accordance with the guidance of the Institutional Animal Care and Use Committee of BSRC Al. Fleming.

For the lung cancer model, mice were anesthetized via i.p. injection with xylazine and ketamine. Cancer cells (2 × 10^5^) resuspended in 50ul DMEM (Gibco) and enriched with 20% growth-factor reduced extracellular matrix (Matrigel, BD Biosciences) were intrapleurally injected in the lung parenchyma of mice using a 29G needle (BD Biosciences). For the metastatic cancer model, mice were injected i.v. (intravenously) with 4 × 10^5^ B16F10 cells in 100μl DMEM medium. Tamoxifen (50mg/kg, Sigma T5648) dissolved in corn oil (Sigma C8267) was given at Col1a2-cre mice i.p. once per day for five consecutive days prior to inoculation of tumor cells.

Monoclonal antibody aCD4/GK1.5 (ATCC/ TIB-207) was administered intraperitonealy (i.p.) 2 days before tumor implantation and continuing 3 times per week for the duration of the study (150ug/mouse). FTY720 (Cayman Chemicals, USA) was administered i.p. for 5 consecutive days, starting on day 7 after tumor cell transplantation at 20mg per mouse.

For the adoptive T cell transfer experiments, splenocytes from OT-II/Rag1/CD45.1 mice were cultured in a-CD28 coated plates (Biolegend, Cat. No. 102115, 1ug/ml) with complete RPMI (10%FBS, 1% Pen/Strep, 50uM b-mercaptoethanol) supplemented with 1ug/ml OVA Peptide (323-339) (Genscript, Cat. No. RP10610) (DAY0). Untouched CD45.1 OTII cells were magnetically sorted from cultured splenocytes on DAY2 using anti-PE MicroBeads (Miltenyi Biotech) after incubation with the following PE-conjugated antibodies: CD11b, CD11c, CD19, B220, CD49b, CD105, MHCII and TER119, according to manufacturer’s instructions. T cells were cultured in the aforementioned medium supplemented with 1ug/ml OVA Peptide (323-339) and 50U/ml mIL-2 (Peprotech, Cat. No. 212-12). On DAY4 OTII cells were transduced with C1qbp lentiviral particles (Lenti ORF, C1qbp, Origene, NM_007573) versus Lenti-ORF Control Particles at a multiplicity of infection (MOI) of 2. The transduction was performed for 24 hours in medium supplemented with 8ug/ml polybrene (Sigma-Aldrich). Two days after lentiviral transduction cells were placed in puromycin selection for 2 weeks (2ug/ml, Sigma P8833). On DAY22 100.000 T cells per mouse were injected intravenously in LLC-OVA tumor-bearing mice.

### RNA extraction and qPCR

For assessment of human C1q expression, total RNA was extracted from apCAFs, nonapCAFs, blood and intratumoral CD4^+^T cells that were isolated as stated above, using Single Cell RNA Purification Kit (Norgen Biotek) according to manufacturer’s instructions. Superscript II reverse-transcriptase (Thermo Fisher Scientific) was used for cDNA synthesis and SYBR Green (Thermo Fisher Scientific) for qPCR performed on the CFX96 Touch™ Real-Time PCR Detection System from Bio-Rad. Transcript levels of C1q were determined relative to U6 reference gene, using the ∆∆Ct method. The following primers sets were used: human C1qb Fw: TAAAAGGAGAGAAAGGGCTTCCAGGG and Rv: TGGCCTTGTAGTCTCCCGATTCACC and human U6, Fw: 5’-CTCGCTTCGGCAGCACA-3’ and Rv: 5’-AACGCTTCACGAATTTGCGT-3’.

### Flow cytometry

FACS analysis or sorting was performed using FACSCANTO II (BD Biosciences) or BD FACSARIA III (BD Biosciences) and data were analyzed using Flowjo software. For calculation of absolute numbers of tumor cells (burden) or immune cells, counting beads (123Count eBeads, ThermoScientific) were used.

### Immunohistochemistry/ Immunofluorescence

For mouse studies, lungs were excised from healthy and tumor bearing mice and fixed in 4% freshly prepared paraformaldehyde for 14-18 hours at 4oC. After fixation and incubation in 30% sucrose, samples were embedded in O.C.T (VWR Cat. 361603E), then sectioned (10μm thick) onto SuperFrost Plus™ microscope slides (Thermo Scientific™). Slides were washed using PBS followed by 1 hour incubation in blocking solution (1% BSA, 0.1% Saponin, PBS). Sections were incubated with primary antibodies Anti-Mouse MHC-II 1:250 (EBiosciences 14-5321-81) and Anti-Mouse Podoplanin eFluor® 660 1:500 (EBiosciences 50-5381-82) diluted in BSA 1%/ PBS overnight at 4oC. After rinsing with 0,1% Saponin/PBS, sections were incubated with secondary antibody for MHC-II (Goat anti-Rat IgG (H+L) Alexa Fluor 594™) for 1 hour at room temperature. Nuclei were stained with DAPI and slides were mounted using Fluoroshield™ mounting media (Sigma F6182). Images were acquired with a TCS SP8X White Light Laser confocal system (Leica).

For IHC and IF stainings of human formalin-fixed paraffin-embedded (FFPE) tissues, 5μm thick sections on charged glass slides were deparaffinized in xylene and rehydrated with ethanol. Sections were incubated for antigen retrieval (Tris EDTA, pH9) for 20 minutes in the microwave and let cool down for 20 minutes at room temperature. For IHC, endogenous peroxidase was blocked by applying UltraVision Hydrogen Peroxide Block (ThermoScientific) for 15 min. Nonspecific protein-binding sites were blocked with UltraVision Protein Block (ThermoScientific) for 5 minutes. Sections were stained with anti-FAP (R&D AF3715) 1:100 or anti-HLA-DR+DP+DQ (Abcam ab7856) 1:200 overnight at 4 oC. Immunodetection was performed using UltraVision Quanto Detection System HRP Polymer DAB (ThermoScientific) according to the manufacturer’s instructions. 3,3′diaminobenzidine Quanto Chromogen (ThermoScientific) was used as chromogen. Slides were counterstained with hematoxylin. For IF, nonspecific protein-binding sites were blocked with 2% BSA/ 2% NGS. Then sections were incubated with anti-FAP (R&D AF3715) 1:100, anti-HLA DR+DP+DQ (Abcam ab7856) 1:200 and anti-CD4 (Abcam ab133616) 1:100 overnight at 4 oC. After rinsing, sections were incubated with secondary antibodies (Goat anti-Rabbit IgG (H+L) Cross-Adsorbed ReadyProbes™ Secondary Antibody, Alexa Fluor 594 Catalog # R37117, Goat anti-Mouse IgG (H+L) Highly Cross-Adsorbed Secondary Antibody, Alexa Fluor 488 Catalog # A-11029 and Donkey anti-Sheep IgG (H+L) Cross-Adsorbed Secondary Antibody, Alexa Fluor 647 Catalog # A-21448). Nuclei were stained with DAPI and slides were mounted using Fluoroshield™ mounting media (Sigma F6182). Images were acquired with a TCS SP8X White Light Laser confocal system (Leica).

Immunofluorescence in T cells for the expression and localization of C1qbp were conducted in sorted intratumoral T cells. After sorting T cells were washed twice and cytospun at 400g for 4 minutes. After cytospin slides let dry for 5-10 min and then cells were fixed for 15 minutes using 4% PFA. After fixation cells were washed twice with PBS 1x and permeabilized for 10 minutes using 0.1% Triton-X/ PBS. Blocking was performed using 2% BSA/ 2% NGS in 0.1% Triton-X 100/ PBS. Cells were stained overnight at 4oC using Recombinant Anti-GC1q R antibody [EPR23238-6] (Abcam ab270033).

### Image acquisition and analysis

Images were acquired using a TCS SP8X confocal system (Leica). We selected a fluorophore panel which allowed for simultaneous visualization of three targets and a nuclear stain (DAPI). During acquisition fluorophores were excited with 405nm (UV Laser), 488 nm (Argon), 594 nm and 647nm (White Light Laser). For images shown in Figure 1 analysis was performed using Leica LASX. Tile scanning was performed in slides stained for pan-MHCII, CD4, FAP and nuclei were stained using DAPI. Autostitching using 10% overlap was followed by analysis. Imaris 9.6 was used for subsequent image manipulations. After creating a colocalization channel between pan-MHCII and FAP all channels were used to define “primary objects” (surfaces - spots) used to analyze the image (distances, cell number, size, XY positioning etc.). Shortest distance calculation and object identification tools were used for data acquisition and image analysis. Cell density analysis was performed by identifying each “object” respectively in fields of 250.000 μm^2. Data exported from Imaris 9.6 including XY location of CD4^+^, MHCII^+^FAP^+^ and MHCII^+^FAP^−^ objects were used for density plot creation using custom R scripts.

### Identification of human MHCII fibroblasts in scRNA sequencing datasets

Processed data were collected from publicly available paired human lung tumour and healthy lung scRNA-seq datasets (E-MTAB-6149 and E-MTAB-6653). The log2cpm values of the 22180 ensembl gene IDs were used. The transition from the raw sequence read data to the gene expression table can be found in the original publication (24). Processed cells annotated as fibroblasts were included in the analysis. Patient #1 was excluded from the analysis because of the small number of Fibroblasts (9 cells). Patient #2 was excluded from the analysis because of the small number of healthy fibroblasts and dendritic cells (5 and 11 cells respectively). Only fibroblasts of patients #3, #4 and #5 were analyzed (n=643, 155 and 486, respectively). To identify MHCII^+^ and MHCII^−^ fibroblasts, cells annotated as cross-presenting dendritic cells in the same dataset were used as positive control for MHCII expression. The 9 MHCII genes (HLA-DRA,HLA-DRB5, HLA-DRB1, HLA-DQA1, HLA-DQA2, HLA-DQB1, HLA-DQB2, HLA-DPA1, HLA-DPB1) were used for re-clustering fibroblasts and dendritic cells with kmeans function of cluster R package with k=3 (tuned for k=3:10), supported by mean Silhouette information scoring, calculated by silhouette function of cluster R package. This resulted in the splitting of the cells into 3 groups (low, middle and high MHCII expression). Fibroblasts of middle and high groups were considered as MHCII^+^ fibroblasts (206 cells), while those of low group as MHCII^−^ fibroblasts. Gene IDs that had zero expression in all fibroblasts were removed from the analysis (16877 IDs were kept). 15487 out of them were successfully translated into gene names using the biomaRt R package. After the characterization of MHCII^+^ and MHCII^−^ fibroblasts, genes with non-zero expression values in more than 25% of each group were kept (1699 genes for apCAFs and 1388 genes for non-apCAFs). The union of the both lists was considered as the list of the genes that are expressed in at least one group. This resulted in 1785 genes.

### Differential expression and pathway analysis of human datasets

To identify the differential expressed genes between the MHCII^+^ and MHCII^−^ we used the FindAllMarkers function of the Seurat R package, giving as input non-scaled read values of the two clusters. We kept cells that had at least 10 features, and removed ribosomal and mitochondrial genes. This resulted to 79 statistical significant up regulated genes (adjusted p-value <0.05, 67 with logFC>1) and 55 statistical significant downregulated (adjusted p-value <0.05, 48 with logFC< −1). The 67 up and the 55 down-regulated were submitted to DAVID for GO term enrichment analysis. DAVID analysis was performed as previously described (Huang et al., 2009) using the οfficial gene names (hg19) as background. Gene ontology using DAVID (6.8) (Huang et al., 2009) was performed for terms classified as biological processes. An enrichment map was created in Cytoscape (v3.7.2) (Shannon et al., 2003) using all significantly enriched GO terms (FDR < 0.1) with a gene set size between 10-1000. Overlap was used as a metric and the cutoff was set to 0.5. Clustering was performed using AutoAnnotate (v1.3.2) (Kucera et al., 2016) with overlap used as edge weight values.

### Bulk mouse RNA sequencing data acquisition, processing and analysis

Intratumoral CD45^+^CD3^+^CD4^+^ T cells were FACS sorted in RL buffer. Total RNA was isolated by using NucleoSpin RNA (Macherey-Nagel). RNA integrity was assessed on an Agilent Bioanalyzer RNA 6000 Pico Chip, library was constructed using NEB Next Ultra RNA Library Prep Kit. The sequencing platform Ion Torrent PROTON and 3′-UTR sequencing strategy was used. Quality of FASTQ files, obtained after Ion Proton sequencing, was assessed using FastQC, following the software recommendations. Quality of FASTQ files was assessed using FastQC, following the software recommendations. Alignment of sequencing reads to the reference genome was performed using the software HISAT2 (version 2.1.0) with the genome reference version mm10. Bam files containing reads that were uniquely aligned were summarized to read counts table using the GenomicRanges package through MetaseqR2 pipeline and default settings for 3’UTR data. The resulting gene counts table was subjected to differential expression analysis (DEA) with metaseqr2 function, using DESeq2 algorithm for the normalization and statistical testing. Genes with less than 5 counts in 75% of the samples were excluded from downstream analysis. DEA thresholds were set for FDR equal to 0.01 and for logFC +-1, returning 365 down-regulated and 19 up-regulated genes. The 365 down-regulated genes were submitted to DAVID for GO term enrichment analysis. DAVID analysis was performed as previously described (Huang et al., 2009) using the official gene names (mm10) as background. Gene ontology using DAVID (6.8) (Huang et al., 2009) was performed for terms classified as biological processes, as described earlier.

MHCII^+^ CAFs and MHCII^−^ CAFs from tumor-bearing mice were FACS sorted as FSC-A^high^CD45^−^CD31^−^EPCAM^−^mCherry^−^PDPL^+^PDGFRa^+^MHCII^+^ and FSC-A^high^CD45^−^CD31^−^EPCAM^−^mCherry^−^PDPL^+^PDGFRa^+^MHCII^−^ cells respectively. Total RNA was isolated by using Single Cell RNA Purification Kit (Norgen Biotek) according to manufacturer’s instructions. RNA integrity was assessed on an Agilent Bioanalyzer RNA 6000 Pico Chip, library was constructed using NEB Next Ultra RNA Library Prep Kit. The sequencing platform Illumina Novaseq 6000 and Pair-end 150 strategy were used. Quality of FASTQ files was assessed using FastQC, following the software recommendations. Alignment of sequencing reads to the reference genome was performed using a two way alignment procedure of initial mapping with STAR (version 2.7.3) and remapping of the unmapped reads with bowtie2 (version 2.3.5.1) software packages with the genome reference version mm10. The raw bam files were summarized to read counts table using the GenomicRanges through MetaseqR2 R package pipeline, and the resulting gene counts table was subjected to differential expression analysis (DEA) using the R package DESeq2 again through MetaseqR2 pipeline. DEA thresholds set for p-value equal to 0.05 and for logFC 1.5 returned 2216 down-regulated and 164 up-regulated genes. The 418 down-regulated genes with logFC >3 were submitted to DAVID for GO term enrichment analysis, as described earlier.

### Gene set enrichment analysis of CAFs for alveolar genes

Gene set enrichment analysis was performed with GSEA (v4.1.0) software. The normalized count table produced by DESeq2 was used for the GSEA according to software recommendations on the standard GSEA run. For human CAFs, the gene set included marker genes of an alveolar cluster that was identified in the same dataset (24). For murine CAFs, we used a gene set that included marker genes of a murine alveolar cluster that was identified in another dataset (54). The rest of the analysis was performed with the default thresholds.

### Cross-species homology analysis

Upregulated genes in MHCII^+^ versus MHCII^−^ fibroblasts were identified independently for each species, as above, and normalized by dividing the average expression of the gene plus a regularization constant (10e-4) by the average of the cluster averages plus a regularization constant (10e-4). After selecting genes with conserved gene symbols, the normalized expression matrices were log-normalized and their correlation calculated by Pearson correlation distance.

### Statistics and reproducibility

For the G-squared tests of independence the g2Test_univariate function of Rfast R package was used. The p-value was calculated only for the upper tail through the pchisq function of the same package. The Pearson correlation values were calculated with cor base R function.

### Visualization

PCA was performed using the prcomp base R function, with centering but no scaling. MDS plot were drawn in R with ggplot2. Box plots were generated using the ggplot2 R package and default parameters. Violin plots were generated using the geom_violin function of ggplot2 R package, with default parameters. MA plots and GO terms dot plot were generated using the geom_point function of ggplot2 R package. Plots were generated using the ggplot2 with dropping. Heatmaps were generated using the pheatmap function of pheatmap R package.

---

**Supplementary Figure 1.**
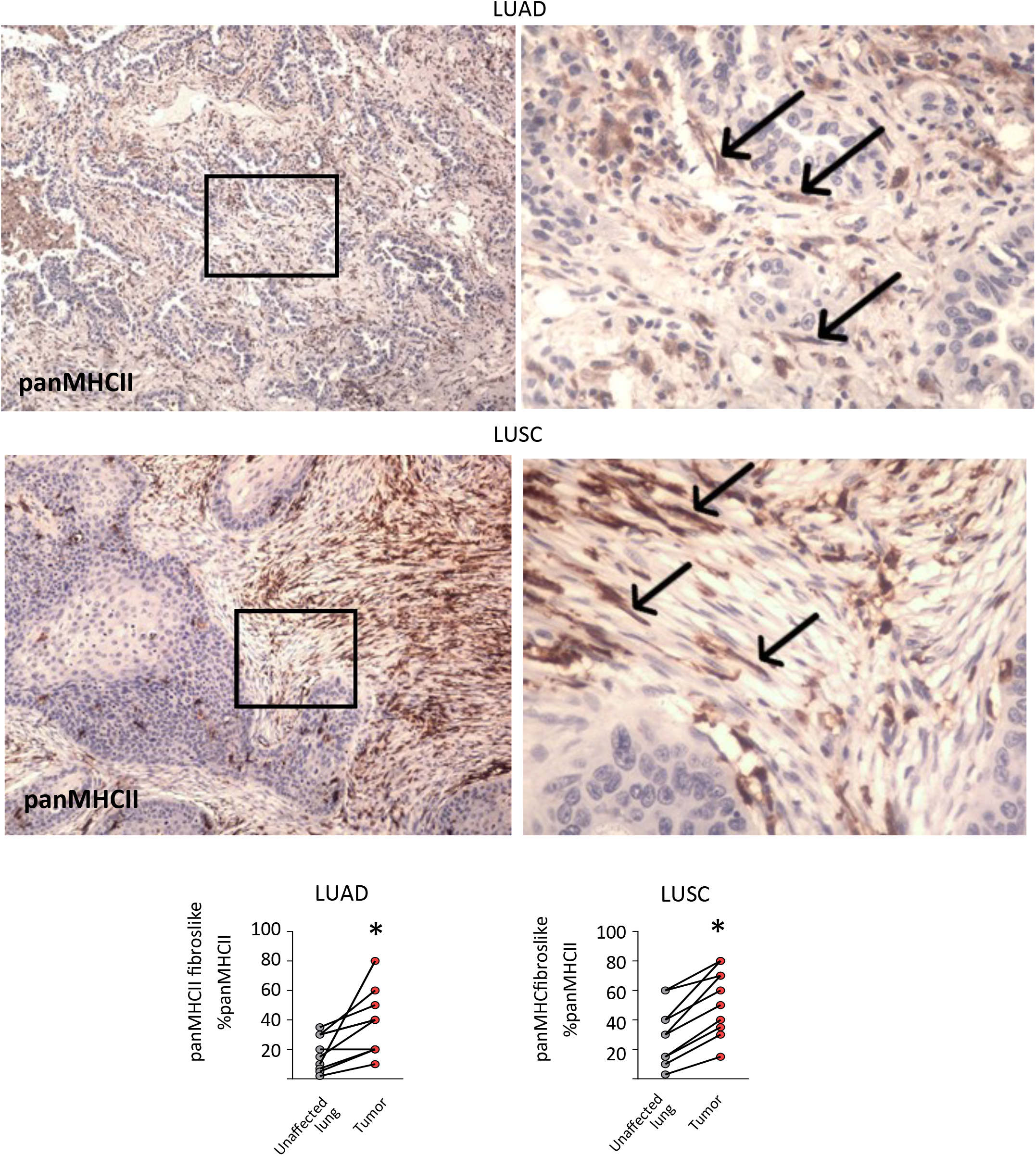
CAFs in human lung adenocarcinomas and squamous cell carcinomas express MHC class II. Representative images (up) and standardized enumeration (down) from immunohistochemistry (IHC) stainings of tumour sections from human lung adenocarcinoma (LUAD) (n=10 patients) and squamous cell carcinoma (LUSC) (n=10 patients), using panMHCII antibodies. Arrows are pointing to examples of CAFs. P<0.05, Paired t-test.

**Supplementary Figure 2.**
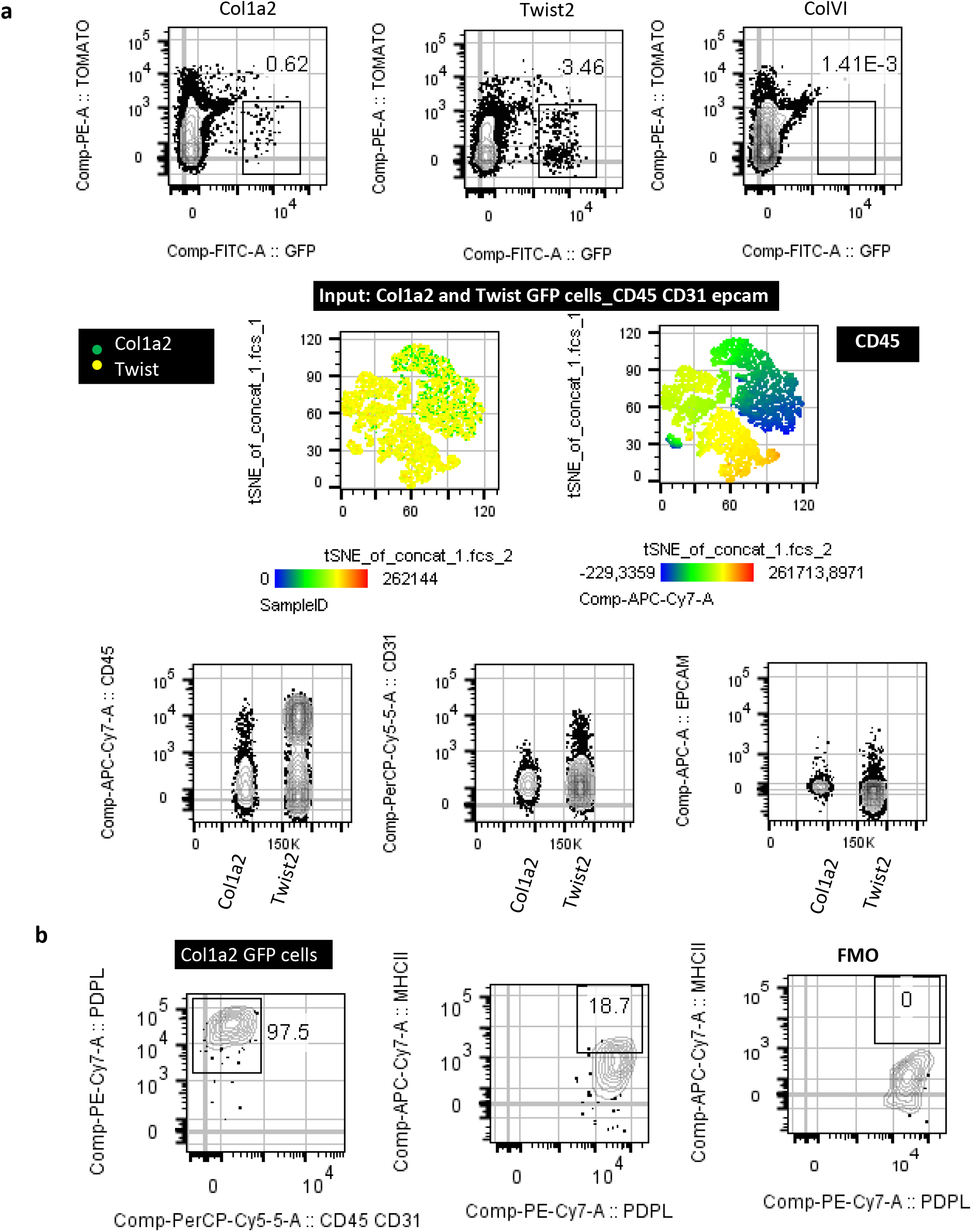
Lug CAF specificity of the Col1a2-CreER, ColVI-Cre and Twist2-Cre mouse strains. LLC cells were implanted intrapulmonary in the left lung lobe of tamoxiphen-treated Col1a2-CreER/mTmG (Col1a2), ColVI-Cre/mTmG (ColVI) and Twist2-Cre/mTmG mice (Twist) mice. Lung tumors were digested and subjected to FACS analyses. **a)** Expression of hematopoietic (CD45), endothelial (CD31) and epithelial (epcam) specific markers by lung GFP cells. **b)** Twist2Expression of the fibroblast-specific marker podoplanin and of IAb (MHCII) (n=2 per group, from 2 independent experiments).

**Supplementary Figure 3.**
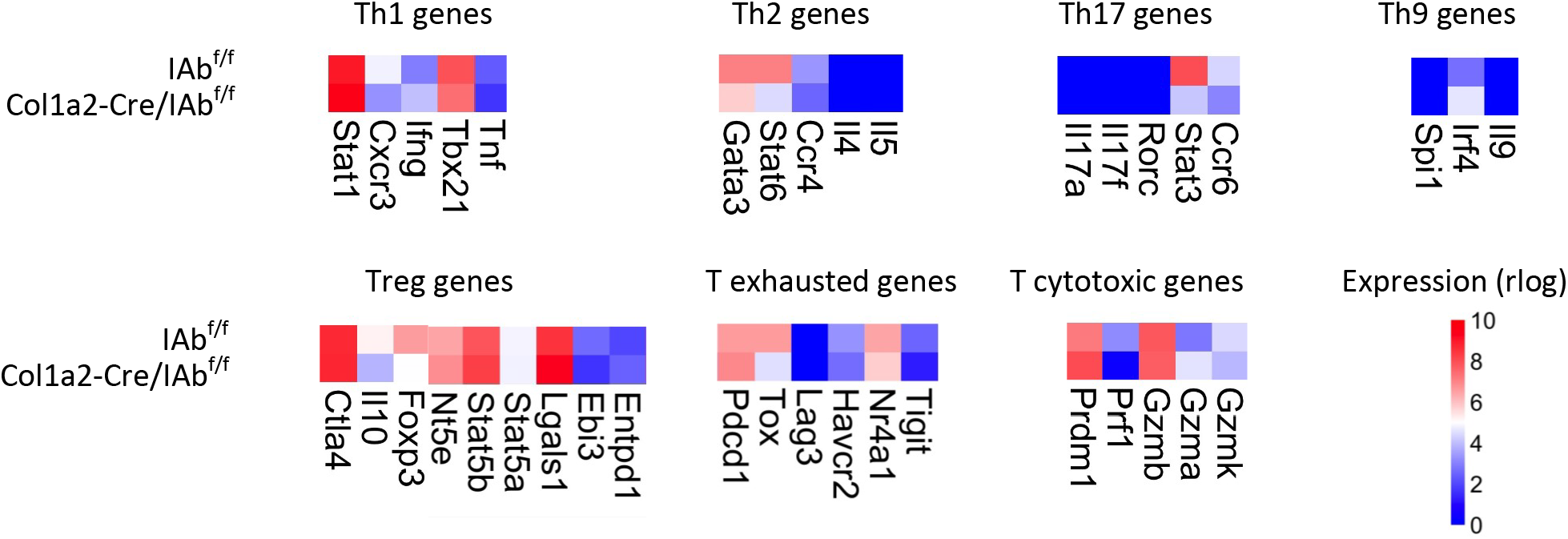
Tumor-infiltrating CD4 T cell marker genes in the Col1a2-CreER strain. Col1a2 CreER^+^I-Ab^fl/fl^ and Col1a2 CreER^−^I-Ab^fl/fl^ mice were inoculated with LLC^mcherryOva^. Differential expression between purified intratumoral CD4 T cells (n=7). Expression heatmaps (mean value) are shown. Adj. P values >0.05, except from: Stat6 0.0042, Stat3:0.0053.

**Supplementary Figure 4.**
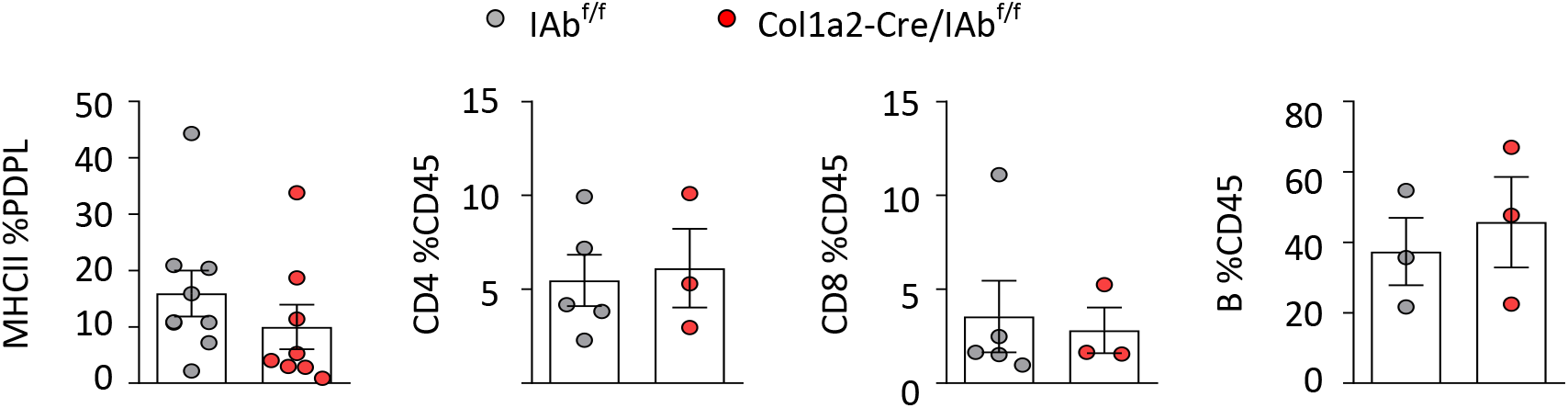
FACS analysis of tumor-draining lymph nodes of the Col1a2-CreER strain. LLC cells were implanted intrapulmonary in the left lung lobe of Col1a2 CreER^+^I-Ab^fl/fl^ and Col1a2 CreER^−^I-Ab^fl/fl^ mice. After 14 days mesothoracic/tumour draining lymph nodes were digested and analyzed by FACS. From left to right. MHCII expression in CD45^−^CD31^−^EPCAM^−^ Podoplanin^+^. Frequencies of CD4^+^, CD8^+^ T cells and B220^+^ B cells. n=3-5 per group, pulled from 2 independent experiments.

**Supplementary Figure 5.**
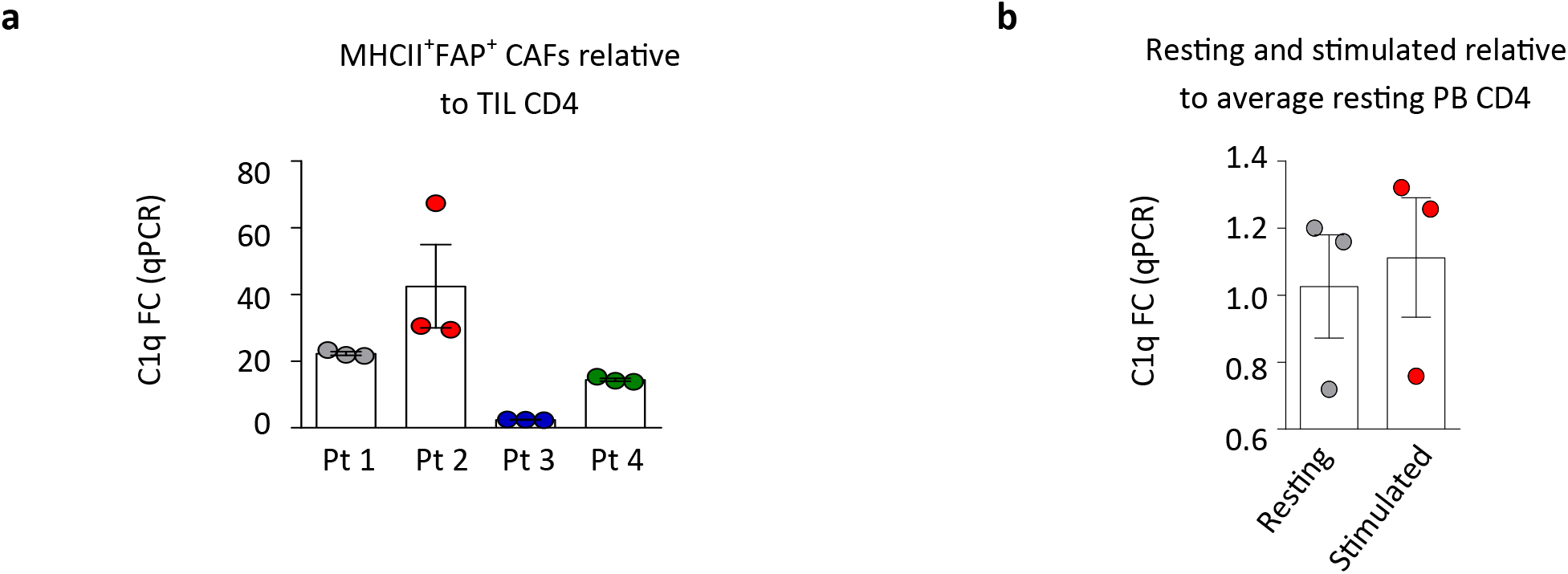
C1q gene expression in primary human apCAFs and CD4 T cells. **a)** Primary human lung tumors were digested. ap-CAFs were sorted as Lin^−^MHCII^+^FAP^+^ cells. Intratumoral CD4 T cells (TIL CD4) were sorted as CD45^+^CD3^+^CD4^+^. RNA was extracted and subjected to qPCR analysis. Relative C1q expression, measured by B-chain component (C1qb). n=4 patients **b)** Untouched peripheral blood CD4 T cells were sorted from a healthy donor and stimulated with aCD3aCD28 beads for 2 weeks. Relative C1q expression, measured by B-chain component (C1qb).

**Supplementary Table 1.** Patient information. Table lists main patient data including number of patients enrolled in the study, age, gender, histology, smoking history and Tumour, Node, Metastasis (TNM) stage.

|  |  | NSCLC |
| --- | --- | --- |
| **Subjects (n)** |  | **32** |
| **Age (years)** |  | **68,71 ± 8,2** |
| **Males** |  | **29 (90,6%)** |
| **Females** |  | **3 (9,4%)** |
| **Smoking Status** |  |  |
|  | Current Smoker | 10 (31,3%) |
|  | Ex -Smoker | 5 (15,6%) |
|  | Unknown | 17 (53,1%) |
| **Tumour Cell Type** |  |  |
|  | Adenocarcinoma | 15 (46,9%) |
|  | Squamous carcinoma | 17 (53,1%) |
| **Overall TNM Stage** |  |  |
|  | IIA | 3 (9,4%) |
|  | IIIA | 7 (21,8%) |
|  | IA2 | 2 (6,3%) |
|  | IA3 | 2 (6,3%) |
|  | IB | 6 (18,7%) |
|  | IIB | 12 (37,5%) |

## Notes

### Competing Interest Statement

The authors have declared no competing interest.

